# Development of a 3D Bioprinted Airway Smooth Muscle Model for Manipulating Structure and Measuring Contraction

**DOI:** 10.1101/2022.12.15.520464

**Authors:** Jeffery O Osagie, Sanjana S Syeda, Emily Turner-Brannen, Michelle Guimond, Lumiere Parrenas, Ahsen Haroon, Philip Imasuen, Adrian R West

**Author notes:** Denotes equal contributions (Jeffery O Osagie, Sanjana S Syeda). **CORRESPONDENCE**, Dr Adrian R West, Children’s Hospital Research Institute of Manitoba, 606A John Buhler Research Centre, 715 McDermot Avenue, Winnipeg MB R3E 3P4, Canada.

## Abstract

The contractile function of airway smooth muscle (ASM) is inextricably linked to its mechanical properties and interaction with the surrounding mechanical environment. As tissue engineering approaches become more commonplace for studying lung biology, the inability to replicate realistic mechanical contexts for ASM will increasingly become a barrier to a fulsome understanding of lung health and disease. To address this knowledge gap, we describe the use of 3D bioprinting technology to generate a novel experimental model of ASM with a wide scope for modulating tissue mechanics.

Using a stiffness modifiable alginate-collagen-fibrinogen bioink, we demonstrate that modulating the stiffness of free-floating ASM ‘bare rings’ is unfeasible; bioink conditions favorable for muscle formation produce structures that rapidly collapse. However, the creation of novel ‘sandwich’ and ‘spiderweb’ designs that encapsulate the ASM bundle within stiff acellular load bearing frames successfully created variable elastic loads opposing tissue collapse and contraction. Sandwich and spiderweb constructs demonstrated realistic actin filament organisation, generated significant baseline tone, and responded appropriately to acetylcholine, potassium chloride and cytochalasin D. Importantly, the two designs feasibly simulate different mechanical contexts within the lung. Specifically, the sandwich was relatively compliant and subject to plastic deformation under high contractile loads, whereas the stiffer spiderweb was more robust and only deformed minimally after repeated maximal contractions.

Thus, our model represents a new paradigm for studying ASM contractile function in a realistic mechanical context. Moreover, it holds significant capacity to study the effects of ECM composition, multiple cell types and fibrosis on lung health and disease.

**GRANTS:**

1. Natural Sciences and Engineering Research Council, Discovery Grant (Adrian West)
2. Research Manitoba, New Investigator Operating Grant (Adrian West)
3. Children’s Hospital Research Institute of Manitoba, Operating Grant (Adrian West)
4. Canadian Foundation for Innovation, John R. Evans Leaders Fund (Adrian West)
5. University of Manitoba, Manitoba Graduate Scholarship (Jeffery Osagie)
6. Research Manitoba, Master’s Studentship Award (Jeffery Osagie)
7. Research Manitoba, Master’s Studentship Award (Sanjana Syeda)
8. Children’s Hospital Research Institute of Manitoba, Summer Studentship (Michelle Guimond)
9. University of Manitoba, Jack Prior Memorial Undergraduate Student Research Award (Lumiere Parrenas)
10. University of Manitoba, Undergraduate Research Award (Ahsen Haroon)
11. University of Manitoba, UMSU Undergraduate Research Award (Philip Imasuen)

The grant bodies had no role in study design, data collection and analysis, decision to publish, or preparation of the manuscript.

## INTRODUCTION

Cell culture, laboratory animals, and *ex vivo* human tissue have been the foundation of biological research for over 100 years. While each remains an essential scientific tool, it is becoming increasingly recognised that these paradigms are not sufficient to fully understand both healthy and disease states. Biological and technical issues limit the tissue- and organ-level relevance of cell culture, while ethical concerns mandate reductions in the use of laboratory animals, and reliable supplies of human tissue remain scarce. Tissue engineering has emerged as a compelling solution to these limitations, by promising to create a large supply of customised human tissues and advanced experimental models from cultured cells to better understand basal biology and to simulate human health and disease (1). In the context of the lung, many recent advancements have been made with ‘organ-on-a-chip’ and bioreactor models that replicate the alveolar compartment, gas exchange, and cell-cell interactions in the airway wall (2–6). Organoid models show extreme promise for modelling different aspects of lung development and epithelial function (7–10).

Despite these advances, major deficiencies persist in our ability to engineer and study a critically important lung component: airway smooth muscle (ASM). ASM composes a large portion of the airway wall by volume, making it a major inflammatory mediator (11). Moreover, muscle contraction is potently modulated by the mechanical environment and accurately replicating these *in vivo* environments *in vitro* has proven to be challenging (12, 13). Tension development and contraction of ASM also regulate vital aspects of lung physiology, and dysfunctional ASM mechanics is a central player in the pathogenesis of lung diseases such as asthma (14). Key technical difficulties arise in the ability to engineer realistic mechanical environments that can stand up to the rigors of muscle contraction. Early attempts successfully used simple hydrogels to measure muscle contraction (15–17), although the mechanical context of these models is questionable. ASM microtissues manufactured on silicone moulds (18), as well as cells on elastomeric substrates (19) and electrospun scaffolds (20), have more realistically replicated many aspects of functional ASM, although short lifespans in culture and narrow scope to modulate tissue mechanics remain significant limitations.

The recent description of a 3D bioprinted experimental model of ASM (21) represents a novel approach for creating functional ASM tissues. In this study, a commercial bioprinter (Aspect Biosystems’ RX1^TM^) and alginate-collagen bioink (AGC-10) were used to create ASM muscle rings that responded appropriately to physiological contractile and relaxant signals. However, similar to early hydrogel approaches, this model does not recreate a realistic mechanical environment, necessitating further development to accurately depict different environments within the lung. Alginate has been used in many tissue engineering applications to modulate tissue stiffness (22–25), and the RX1 bioprinter incorporates a multi-channel microfluidic printhead that could feasibly deposit structural and muscle components separately, presenting several avenues for technical and physiological advancement.

In the present study, we expanded on previous approaches by creating a stiffness-modifiable alginate bioink and novel multi-material tissue structures. This enabled us to develop a 3D bioprinted model of ASM designed specifically to interrogate the effect of tissue mechanics on muscle contraction. Critically, our approach generated lifelike ASM that responded appropriately to contractile and relaxant signals and demonstrated that different engineered structural properties can modulate the dynamics of ASM contraction. Finally, by testing coronary artery smooth muscle (CSM) in parallel, we demonstrated that our model could be applicable to a wide range of different organ contexts.

## METHODS

### Cell Culture

Senescence resistant human ASM cells (hTERT overexpression, line D12) were obtained as a generous gift from the laboratory of Dr. Andrew Halayko (Department of Physiology & Pathophysiology, University of Manitoba). Cells were propagated in feeder media consisting of DMEM/F12 (Gibco 11330-057) supplemented with 10% heat-inactivated fetal bovine serum (Gibco 12483-020) and 1% penicillin-streptomycin (Gibco 15140-122), and maintained at 37°C, 5% CO_2_ and 85% humidity. To prepare for bioprinting, cells at approximately 90% confluence were detached with TrypLE cell dissociation reagent (Gibco 12605-036), resuspended in a calcium-free alginate diluent buffer (145 mM NaCl, 5 mM NaHCO3, pH 7.0), and gently centrifuged at 150×RCF for 5 minutes. The supernatant was aspirated, and cells resuspended into prepared bio-ink at a density of 2.5×10^7^ cells/mL. Coronary Arterial Smooth Muscle cells (CSM; Applied Biological Materials T0557) were propagated and prepared in the same manner, but were grown in flasks coated with rat tail collagen type I (Corning 354236) at 10 μg/mL in PBS.

### 3D Designs

Designs for 3D bioprinting and fused deposition modelling (FDM) 3D printing were created in Tinkercad (https://www.tinkercad.com/) and exported as STL files (see Supplementary Files). Toolpaths for 3D bioprinting were prepared from STL files in Aspect Studio software and saved as APJ files. ‘Bare ring’ designs were created as a cylinder 8 or 10 mm in diameter and 0.6 mm high and printed with 0.04 mm layer thickness (for 15 layers), 0% infill, and one perimeter of cellular bioink (Fig 1A). ‘Sandwich’ designs encapsulated a bare ring within top and bottom 13 mm diameter, 0.20 mm high cylinders as acellular layers (Fig 1B). The encapsulation layers were set to a 0.04 mm layer height (for 5 layers), and 20% infill with rectilinear and concentric patterns in alternate layers. ‘Spiderweb’ designs were created with additional mechanical supports designed to interface with a rigid plastic tissue holder (Fig 1C). Briefly, tissue holders were designed with six internal posts and an outer diameter set to fit securely inside a 24 mm ThinCert insert (Greiner 657638). We hand-coded bioprinter toolpaths to generate a 3D design comprising a 10-layer 10 mm diameter cellular ring stacked between four layers of acellular rectilinear infill from the sandwich design. This was further encapsulated by a 19 mm diameter network of acellular material containing loops/voids positioned to engage with the tissue holder. Layer heights were set such that the spiderweb construct was nominally 1.0 mm thick. To improve stability of constructs for long-term live cell imaging, centring rings were designed to firmly hold the ThinCert inserts at the centre of a 6-well microplate (Greiner 657160) without disrupting access to the basolateral chamber (see Supplementary Files).

**Figure 1.**
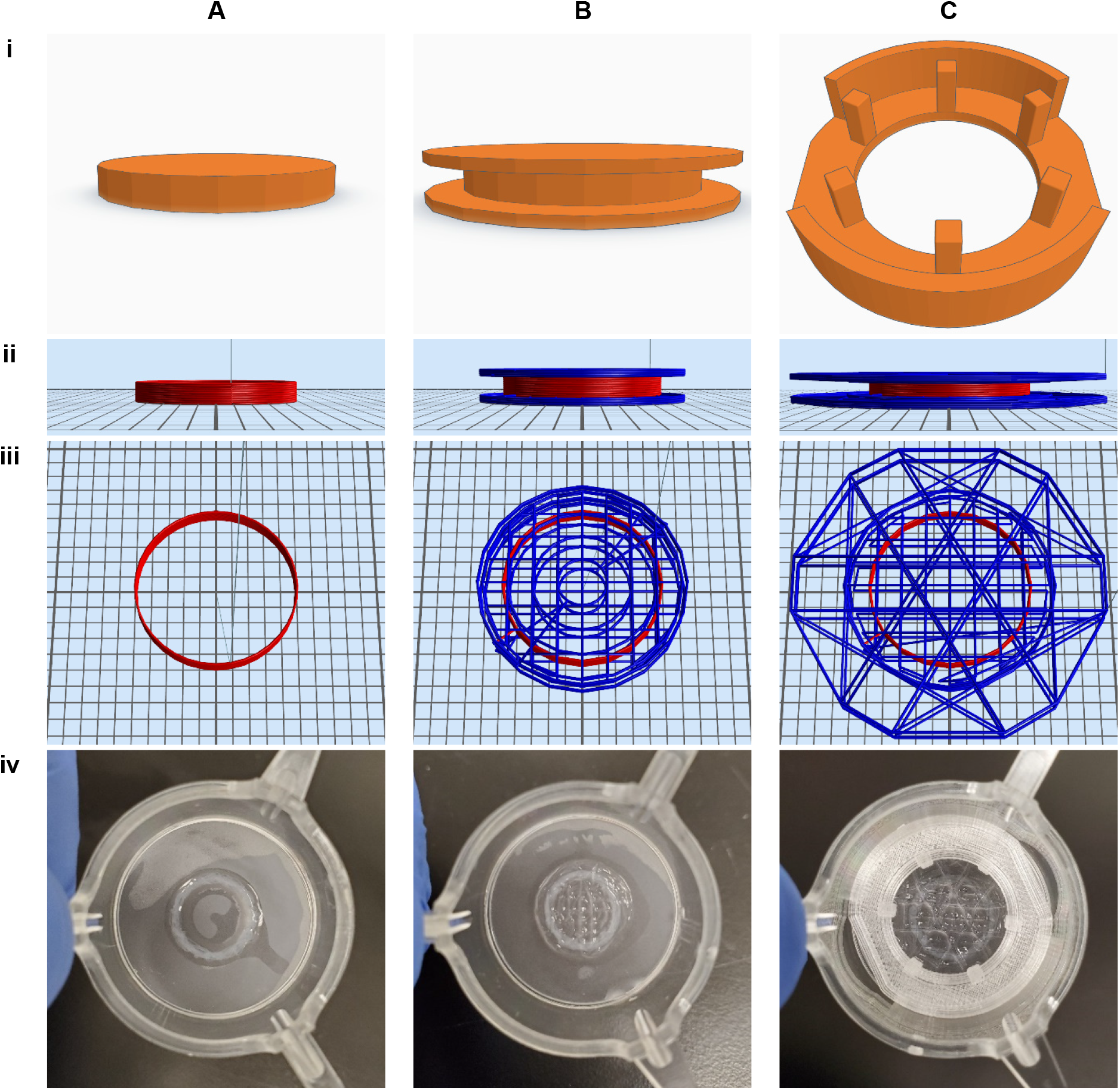
3D STL designs and toolpaths for bioprinting and FDM printing. Schematic of bare rings (A), sandwich (B), and spiderweb (C) designs showing STL files (i), bioprinter toolpaths (ii and iii) and photos of freshly printed constructs (iv). Bare rings consisted of a 10 mm diameter cylinder printed with cellular bioink (red) in a single circumference 15 layers high. The sandwich design encapsulated the bare cellular ring between two 13 mm dimeter cylinders, with alternating rectilinear and circular infill patterns of acellular bioink (blue) creating a spring-like load opposing tissue compaction and contraction. The spiderweb design began as a solid tissue holder; the posts were designed to engage with voids in a hand-coded toolpath for acellular bioink that further encapsulated and supported the cellular ring. Each of the three are printed with very high fidelity to the base design. The Z-height of ring and sandwich STL files, plus all toolpaths, are scaled to 200% for visual clarity.

### FDM 3D Printing

STL files were 3D printed using Ultimaker Cura 4 slicing software and an Ultimaker S3 3D printer (216932) fitted with an AA 0.25 mm print core (200454), glass print surface (227639) and lid enclosure (Shop3D EnclosureS3). Filament was stored in freshly regenerated silica gel between prints and kept within a PrintDry 2.0 filament dryer at ≥65°C during printing to minimise moisture content and ensure print consistency. Tissue holders were printed with Polycarbonate Transparent filament (Ultimaker 1640) using a modified version of the Cura Fine (Experimental) profile (see Supplementary Files) and an adherence aid to prevent warping (Magigoo PC). Printing the fine features of the tissue holder exceeded filament retraction limits, thus sacrificial cylinders were interspersed between the tissue holders to consume filament and enable unlimited retractions without filament damage (see Supplementary Files). Completed tissue holders were visually inspected, and those with excessive stringing or obvious physical defects were discarded. Acceptable prints were washed several times with distilled water then once with 70% EtOH to remove traces of the adherence aid, air dried, then passed briefly through a Bunsen burner flame to remove minor surface stringing. Tissue holders were sanitized by boiling in ddH2O water for 10 minutes at full power in a 1000 W microwave oven, followed by soaking in 70% EtOH under moderate vacuum for 30 minutes. They were then exposed under 70% EtOH to 400 mJ/cm^2^ of UV-C light per side in a Spectrolinker XL-1000 UV Crosslinker, before being transferred to a sterile field for air drying and storage. ThinCert centring rings were printed with Ultrafuse PET Black filament (BASF Pet-0301a075) using the Cura CPE Generic Fine profile with modifications (see Supplementary Files) and Magigoo Original adherence aid. Centring rings were sanitised by the same process as for tissue holders, with the boiling step omitted.

### Bioink Formulation

All bioinks were comprised of alginate, collagen, and fibrinogen as appropriate to individual experiments. To create stiff bioinks, we used an ultra-purified high guluronate low molecular weight alginate (NovaMatrix PRONOVA SLG-20; referred to as SLG-20). Biocompatibility and soft bioinks were achieved with an RGD-coupled high guluronate very low molecular weight alginate (NovaMatrix NOVATACH VLVG GRGDSP; referred to as RGD-VLVG). Lyophilised alginates were diluted to working concentration of 3% w/v by adding alginate diluent buffer and incubating at room temperature with gentle mixing until fully dissolved. Fibrinogen (Sigma F8630-1G) was freshly prepared as a 20 mg/mL stock solution by carefully layering the powder on top of alginate diluent pre-warmed to 37°C. The tube was gently swirled to mix and maintained at 37°C until fully dissolved before the solution was sterilized by passing through a syringe filter (Sigma SLGV033RS), then kept on ice. Collagen I was added to a pre-cooled 15 mL tube, before pre-calculated volumes of ice-cold 7.5% NaHCO3, 1.5 M NaCl and ddH_2_O were added dropwise while gently vortexing the tube to precisely neutralize the acid from collagen and achieve salt concentrations of 145 mM NaCl, 5 mM NaHCO3 within the final volume of the completed bioink. The appropriate quantities of fibrinogen and alginate were then added dropwise with gentle mixing on ice. A pH strip (5–8) was used to verify that the pH of the final mixture was in the 7.0 to 7.4 range.

### Bioprinter Setup and Printing Process

An Aspect Biosystems RX1 bioprinter was set up according to the manufacturer’s instructions. In brief, a DUO (two bioink material channels) or QUAD (four bioink material channels) microfluidic printhead was mounted on the bioprinter. Material tubes (Falcon 352096) containing bioinks and bottles containing alginate diluent buffer and CAT-2 crosslinker (Aspect Biosystems 00000458) were sealed with pneumatic caps pre-threaded with autoclaved PTFE microfluidic tubing (Diba Omnifit 008T16-080-20), before the material tubes were placed in an ice filled tube holder. Tubing was connected to the corresponding material channels in the printhead, before the system was primed to remove air and clear bubbles from the printhead and nozzle in the following order: material 1 through material 4, buffer, buffer plus crosslinker, then buffer.

ThinCert inserts were pre-wetted with alginate diluent buffer and mounted on the RX1 24 mm vacuum chuck. The porous membrane captured the printed material while allowing excess crosslinker and buffer to flow through. Constructs were deposited layer by layer as per the specified APJ file, with pressures and speeds set and constantly monitored to maintain high fidelity to the original design (Table 1 and Fig 1). All print runs were completed within 30 minutes of adding cells to bio-inks to minimize variance arising from cell sedimentation/clumping and premature precipitation/gelation of bioink components. For spiderweb constructs, a sanitized tissue holder was manually applied immediately after printing to engage the posts with loops/voids in the construct. ThinCert inserts containing the printed constructs were transferred to a 6-well plate containing 4 mL of pre-warmed feeder media. Where the bioink contained fibrinogen, the feeder media was supplemented with 1.25 U/mL thrombin (Sigma T4648-1KU) for 30 minutes to initiate fibrinogen cleavage and polymerisation to fibrin, before constructs were switched to regular feeder media. For spiderweb designs, a ThinCert Centring Ring was applied to prepare the constructs for post-print imaging (see below). Constructs were maintained by changing media every two days. Prior to contraction assays, constructs were serum-deprived for ≥24 hours by replacing feeder media with DMEM/F12 containing 0.5% FBS and 1% insulin-transferrin-selenium (Gibco 41400-045).

**Table 1.** Print pressures and speeds for select bioink formulations during bare ring and sandwich/spiderweb optimisation and experimentation phases. Pressures were qualitatively monitored during printing to maintain fidelity to the original 3D design and comparable deposition rates between different bioink viscosities.

| **Bare ring optimisation** | **Print Pressure (mBar)** | **Print Speed (mm/sec)** |
| --- | --- | --- |
| Cellular 0.25% RGD-VLVG | 18-22 | 32 |
| Cellular 0.375% RGD-VLVG | 20-28 | 32 |
| Cellular 0.5% RGD-VLVG | 25-30 | 32 |
| Cellular 0.75% RGD-VLVG | 30-50 | 30 |
| Cellular 0.75% SLG20 | 55-60 | 23 |
| Cellular 1.0% SLG20 | 80-100 | 20 |
| **Optimised for sandwiches and spiderwebs** | **Print Pressure (mBar)** | **Print Speed (mm/sec)** |
| Cellular, 0.375% RGD-VLVG | 25-35 | 32 |
| Acellular, 0.75% SLG20 | 80-100 | 23 |
| Acellular, 1.0% SLG20 | 120-150 | 20 |
| Acellular, 1.25% SLG20 | 160-200 | 18 |
| Crosslinker | 60-70 | N/A |
| Buffer | 100 | N/A |

### Imaging for Tissue Compaction and Contractile Function

To assess compaction of bare rings and sandwich constructs, macroscopic images were captured daily using an Olympus Tough TG-1 iHS digital camera mounted on a tripod. Camera height and focus were optimized using 6-well plates with ThinCert inserts such that the entire plate was in the field of view. Videos of muscle responses to contractile/relaxant agents and alginate decrosslinking were obtained using high-quality video mode (1080p @ 30 fps). Videos were processed by ffmpegTool to create single image sequences. Spiderweb constructs were imaged in a Cytation 5 MPV Cell Imaging Multimode Reader set at 37°C and 5% CO_2_. Whole-well phase contrast images were obtained using the montage function of the Gen5 software and a 4× lens.

### Contraction Assays

To determine contractile responses of sandwich and spiderweb constructs, time-lapse images representing baseline and treatment conditions were captured. Drug treatments designed to elicit maximal responses were prepared in low serum media; 100 μM acetylcholine (ACh; Sigma A6625), isotonic 80 mM KCl (Sigma, 746436-500) and 10 μM cytochalasin D (Cyto D; Santa Cruz, sc-201442) as described previously (18). To initiate the experiment, the media on constructs was replaced by the first drug treatment, adding one-third of the dose to the ThinCert apical chamber and two-thirds to the basolateral chamber, taking care not to disturb the constructs. After a 10-minute exposure window, treatment images were captured, before applying the next treatment in the sequence. For single maximal contraction time course experiments in spiderweb constructs, a 20-minute KCl exposure was followed by a 20-minute cytochalasin D exposure, with images captured continuously at the maximum capture rate.

### Alginate Depolymerization Load Removal Assay

To characterize shortening dynamics of the unloaded muscle bundle and determine the relative strength of the mechanical load, calcium chelation was used to rapidly de-crosslink alginate (26, 27). Baseline images were taken before media was replaced with a buffer containing 40 mM tri-sodium citrate (Sigma 111037) plus 15 mM ethylenediaminetetraacetic acid (EDTA; Sigma E5134-50G) pH 7.2. Videos (sandwiches) and montage image sequences (spiderwebs) were collected for up to 10 minutes.

### Contraction Reproducibility

To determine the robustness and stability of our designs, spiderweb constructs were subjected to five successive contraction/relaxation cycles. After capturing baseline images, maximal contraction was achieved by 20-minute exposures to 80 mM KCl, followed by a 30-minute washout period, with images captured at peak contraction and relaxation. The final washout cycle was followed by an additional treatment with 10 mM cytochalasin D to ensure complete tension ablation.

### Image Processing and Calculations

The lumen area of bare ring, sandwich, and spiderweb constructs was quantified with Fiji Imaging software, using the manual polygon selection feature. Where suitable, the Canny Edge Detector plugin was used to highlight borders to simplify and expedite measurements. Changes in lumen area were calculated as a percentage of raw pixel area relative to, or delta from, post-print area (for compaction assays) and pre-contraction baselines (for contraction assays).

### Histology

Constructs were fixed with 4% paraformaldehyde (PFA; Sigma P6148) in Hanks Balanced Salt Solution (HBSS; Gibco 14065-056) for 15 min, permeabilized with 0.1% Triton-X 100 (X-100, Sigma-Aldrich) in 4% PFA/HBSS for 10 min, then washed 3 times with HBSS. Filamentous actin (f-actin) was stained using 1 U per construct of Alexa Fluor 488 Phalloidin (Invitrogen A12379) in HBSS for 1 hour, before washing 3 times with HBSS, and counterstained with 0.1 μg/mL Hoechst 33342 (Invitrogen H1399) in HBSS for up to 20 min. Constructs were washed one final time to remove unbound stain before they were transferred to a glass bottomed 12 well plate (Cellvis P12-1.5H-N). One-to-two drops of SlowFade Diamond Antifade Mountant (Invitrogen S36963) were added, and an 18 mm round coverslip (VWR CA48382-041) placed on top. Widefield fluorescence images were captured on the Cytation 5; where necessary, images were uniformly adjusted for brightness and contrast only, ensuring that no details were obscured.

### Data Analysis and Statistics

All numerical data are presented as mean ± standard error. Each ‘n’ value represents results from a single distinct bioprinted construct. Statistical tests were performed with the GraphPad Prism 6.07 software package, with p<0.05 considered statistically significant. For all compaction time-course and contraction experiments, independent samples 2-way ANOVA (factors: Stiffness, Time) was performed, with all-pairwise Tukey post-tests performed for the Stiffness factor. Contractile reproducibility was assessed by independent samples 2-way ANOVA (factors: Contraction/Relaxation, Cycles) with Tukey post-tests comparing contraction versus relaxation within each cycle.

## RESULTS

### Bare Ring Optimisation

To begin formulating a stiffness-modifiable bioink suitable for use with ASM, we took inspiration from Aspect Biosystems’ previous work (21) and began producing bare ring designs containing ASM and CSM with variable concentrations of SLG-20 alginate plus collagen. Rings with high concentrations of SLG-20 were mechanically robust but exhibited no characteristics of functional muscle. F-actin staining revealed that cells within bare rings were balled-up immediately after printing, as would be expected from the dissociation step during bioink preparation (Fig 2A). However, the cells remained balled-up even after extended periods in culture (up to 7 days). As SLG-20 concentration was reduced to produce softer rings, or replaced in-part or completely by the less-stiff RGD-VLVG, cells began to exhibit some signs of cell spreading and elongation. However, these less-stiff alginate formulations resulted in rings that were extremely fragile, collapsed into a shapeless mass of tissue, and began to exhibit tears in the macroscopic structure (Fig 2B).

**Figure 2.**
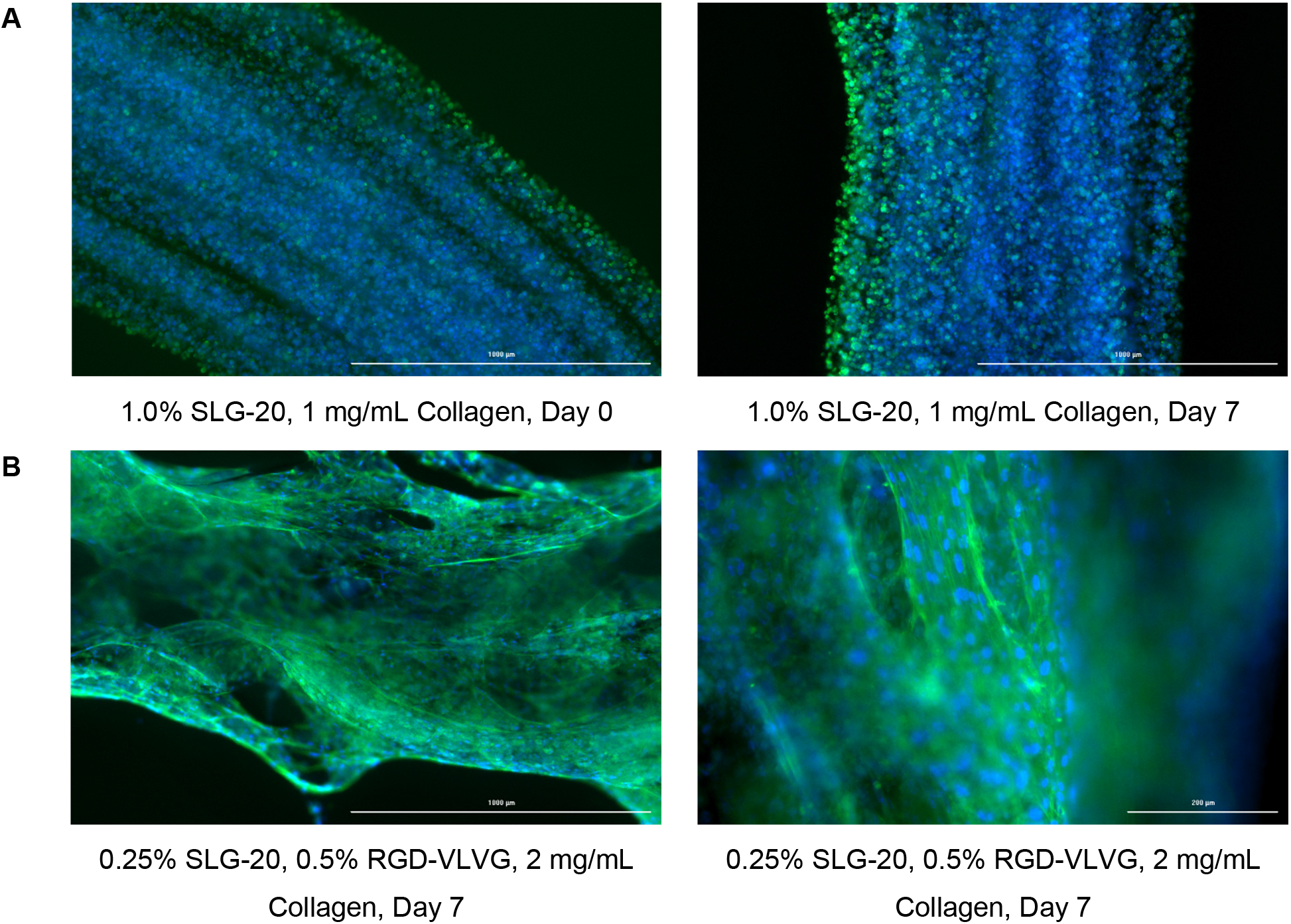
Histology of bare ASM rings with different bioink formulations. Bare rings formulated with high concentrations of SLG-20 produced high fidelity and robust structures (A), however cells did not form functional muscle due to an absence of cell spreading, as evidenced by disorganised f-actin (green) and nuclei (blue) staining even after extended culture periods. Lower concentrations of SLG-20 and substitution of SLG-20 with RGD-VLVG allowed cells to spread within the construct (B). However, the macroscopic structure began to show tears in the fibre and collapses soon after printing. These results were consistent regardless of cell line/type and collagen concentration.

To ensure that this phenomenon was not specific to a particular cell type, several different hTERT ASM cell lines plus CSM were trialled, each with similar findings (data not shown). Increasing collagen concentration also failed to improve either cell spreading or structural integrity in any alginate formulation. This created a significant impasse for optimisation of bare rings; stiff bioinks that facilitated structural integrity did not facilitate muscle formation/maturation, and soft bioinks that supported muscle formation were unable to withstand long-term cell culture conditions due to rapid loss of structural integrity.

### Sandwich Development

Several approaches were trialled to achieve a delicate balance between cell spreading and structural stability in constructs made with low stiffness alginates. Initial attempts to increase internal compressive loads by modifying the aspect ratio of bare rings to make them wider and flatter yielded no benefits. Printing strategies were then developed to exploit the multi-material capabilities of the RX1 bioprinter, by including acellular components comprised of stiff alginate, intended to provide physical support. First, a heterogenous ‘Janus’ fibre design was trialled where cellular and acellular materials were co-deposited into a single fibre, however this proved extremely challenging to print and produced muscle tissue with little functionality. Second, an ‘alternating ring’ approach interspersed stiff acellular layers within the bare ring, yielding minor but insufficient structural improvements (data not shown). Finally, several methods of fully encapsulating the muscle ring within an acellular structure were tested and iterated, ultimately resulting in the sandwich design (Fig 1B). The intent of this design was to reduce mechanical stress/burden on the muscle bundle, allowing the cellular bioink to be formulated with low alginate concentrations that facilitated tissue remodelling, and the acellular bioink could be made with higher and tunable stiffnesses to provide varying loads opposing ring collapse.

An extensive iterative design process was implemented to formulate optimal cellular and acellular bioinks. These were based on SLG-20, RGD-VLVG, collagen I and fibrinogen (Table 2), using a single ASM cell line (D12) and CSM cells to minimise variation, with success measured by qualitative tissue/print fidelity, cell spreading, actin morphology, and tissue lifespan in culture. Notably, inclusion of SLG-20 in the cellular bioink at any concentration completely inhibited muscle formation, and the ideal concentration of RGD-VLVG was relatively narrow. Use of collagen and fibrinogen enhanced both muscle formation and technical aspects of printing, although the exact concentrations of these components did not seem to be critical. Thus, the optimal cellular bioink which offered a balance between practicality and tissue formation contained 0.375% RGD-VLVG, 1.0 mg/mL collagen, 5.0 mg/mL fibrinogen. The optimal acellular bioink contained 0.75-1.25% SLG-20, 1.0 mg/mL collagen, 5.0 mg/mL fibrinogen.

**Table 2.**
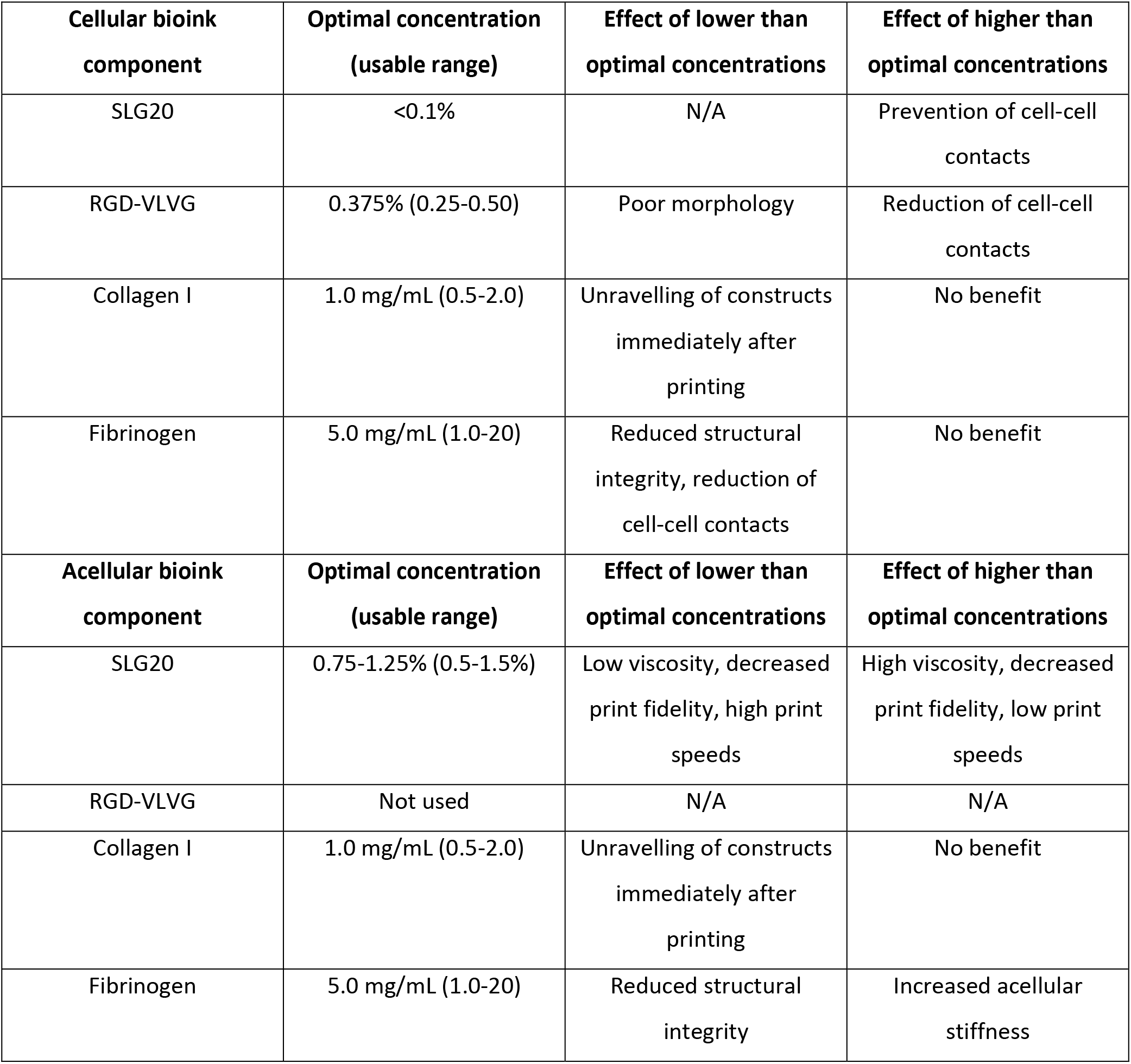
Summary of bioink components and their usable concentration range within cellular (muscle) and acellular (structural) components of 3D bioprinted sandwich and spiderweb constructs. For convenience, the acellular bioink was typically made with the same collagen and fibrinogen content as the cellular bioink.

| **Cellular bioink component** | **Optimal concentration (usable range)** | **Effect of lower than optimal concentrations** | **Effect of higher than optimal concentrations** |
| --- | --- | --- | --- |
| SLG20 | <0.1% | N/A | Prevention of cell-cell contacts |
| RGD-VLVG | 0.375% (0.25-0.50) | Poor morphology | Reduction of cell-cell contacts |
| Collagen I | 1.0 mg/mL (0.5-2.0) | Unravelling of constructs immediately after printing | No benefit |
| Fibrinogen | 5.0 mg/mL (1.0-20) | Reduced structural integrity, reduction of cell-cell contacts | No benefit |
| **Acellular bioink component** | **Optimal concentration (usable range)** | **Effect of lower than optimal concentrations** | **Effect of higher than optimal concentrations** |
| SLG20 | 0.75-1.25% (0.5-1.5%) | Low viscosity, decreased print fidelity, high print speeds | High viscosity, decreased print fidelity, low print speeds |
| RGD-VLVG | Not used | N/A | N/A |
| Collagen I | 1.0 mg/mL (0.5-2.0) | Unravelling of constructs immediately after printing | No benefit |
| Fibrinogen | 5.0 mg/mL (1.0-20) | Reduced structural integrity | Increased acellular stiffness |

Gross morphology of constructs printed with different stiffness acellular bioinks revealed that ASM sandwiches (Fig 3A-B) have modest but significant unstimulated ‘compaction’ of lumen area over time (p<0.0001), indicative of baseline tone development and tissue remodelling. This effect is stiffness modulated (p<0.0001) with 0.75% alginate constructs experiencing significantly higher compaction. Consistent with previous results, ASM bare rings compacted excessively, collapsed, and rapidly became non-functional. F-actin staining for ASM (Fig 3C-D) tissues printed with optimal acellular and cellular bioinks demonstrated impressive cell spreading/elongation and alignment, consistent with native muscle tissue. CSM sandwiches (Fig 3E-F) followed a similar dynamic over time (p<0.0001), but we were unable to detect statistically significant differences in compaction across the stiffness range trialled (p=0.4693). Collectively, these results show that the sandwich design successfully stabilises tissue structure and facilitates lifelike muscle morphology, establishing it as a suitable platform for the assessment of contractile function.

**Figure 3.**
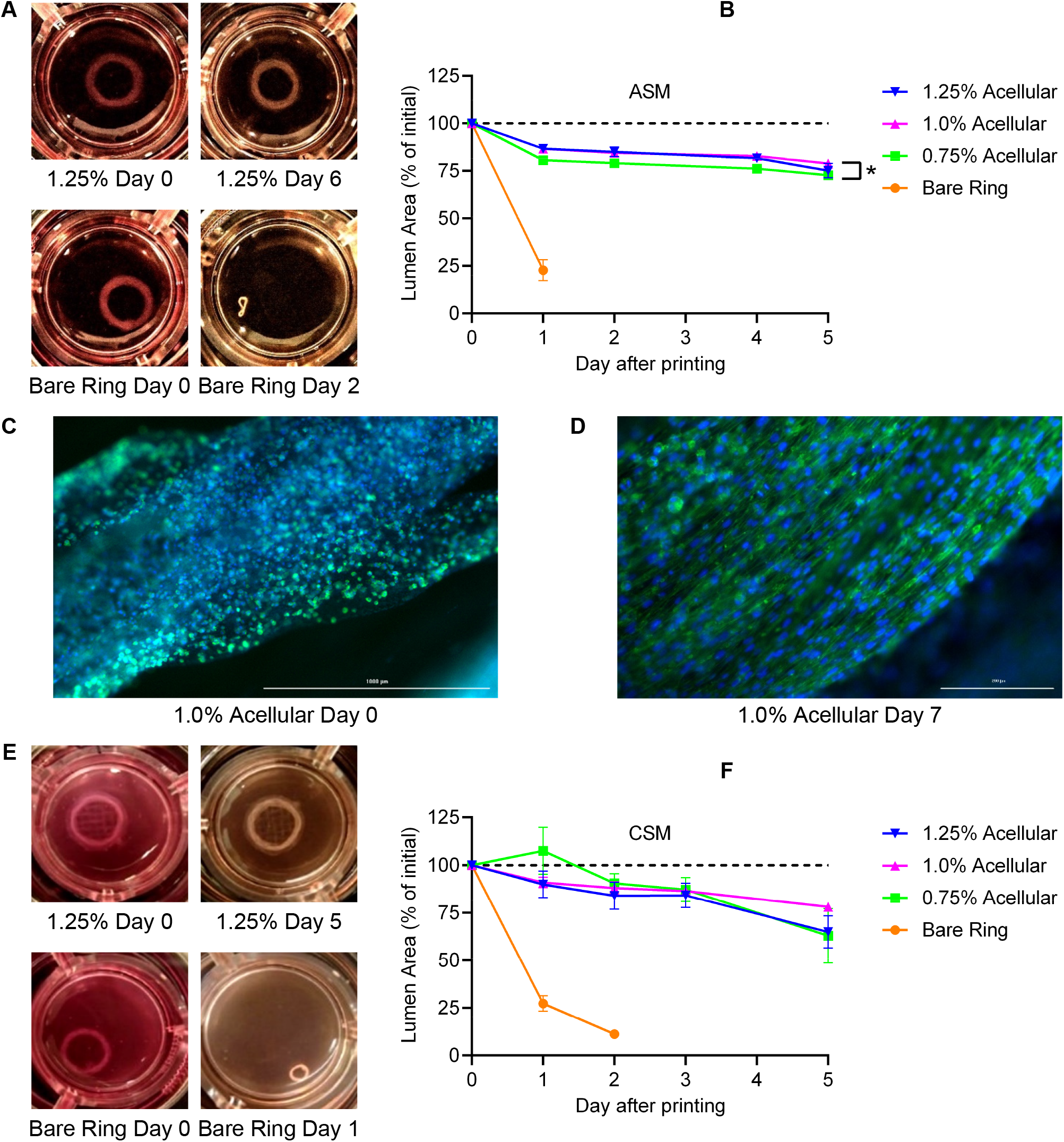
The sandwich design opposes muscle ring collapse and promotes muscle-like morphology. ASM sandwiches are qualitatively equivalent to bare rings immediately after printing, but the sandwich structure effectively prevents ring collapse (A). Compaction of ASM sandwiches is moderately but significantly regulated by times and acellular stiffness (B; interaction p=0.5164, time p<0.0001, stiffness p<0.0001, n=12). Widefield fluorescence of ASM sandwiches for f-actin (green) and nuclei (blue) reveals disorganised and rounded cells immediately after printing (C), but cells rapidly take on an elongated morphology with f-actin alignment along the length of the muscle bundle which is highly reminiscent of real muscle (D). CSM sandwiches are similarly resistant to collapse over extended culture periods (E); they exhibit a highly similar time-based compaction, although this is not modulated by acellular stiffness (F; interaction p=0.5949, time p<0.0001, stiffness p=0.4693, n=4). Note that bare rings are shown on graphs for reference but are excluded from statistical analysis. * represents a significant stiffness effect.

### Sandwich Contractile Function

ASM sandwich constructs responded appropriately to classic contractile stimuli (Fig 4A; p<0.0001), exhibiting mild contractions in response to ACh and a stronger response to KCl. Cytochalasin D reduced lumen area consistent with cytoskeletal tension ablation, however lumen area failed to return to baseline levels. Responses were stiffness modulated (p=0.0438), although this modest effect was not linear, with 1.0% acellular sandwiches having a weaker contractile response than 0.75 and 1.25% constructs. CSM sandwich constructs exhibited similar contractile trends with apparent reductions in lumen area in response to KCl and reversal by cytochalasin D, but the treatment effects were not statistically significant over time due to extremely high variability between tissues (Fig 4B; p=0.1788). There appeared to be a trend towards linear stiffness modulation with 1.25% acellular constructs producing the highest mean contraction and weakest mean relaxation response, but the trend was not significant (p=0.2726).

**Figure 4.**
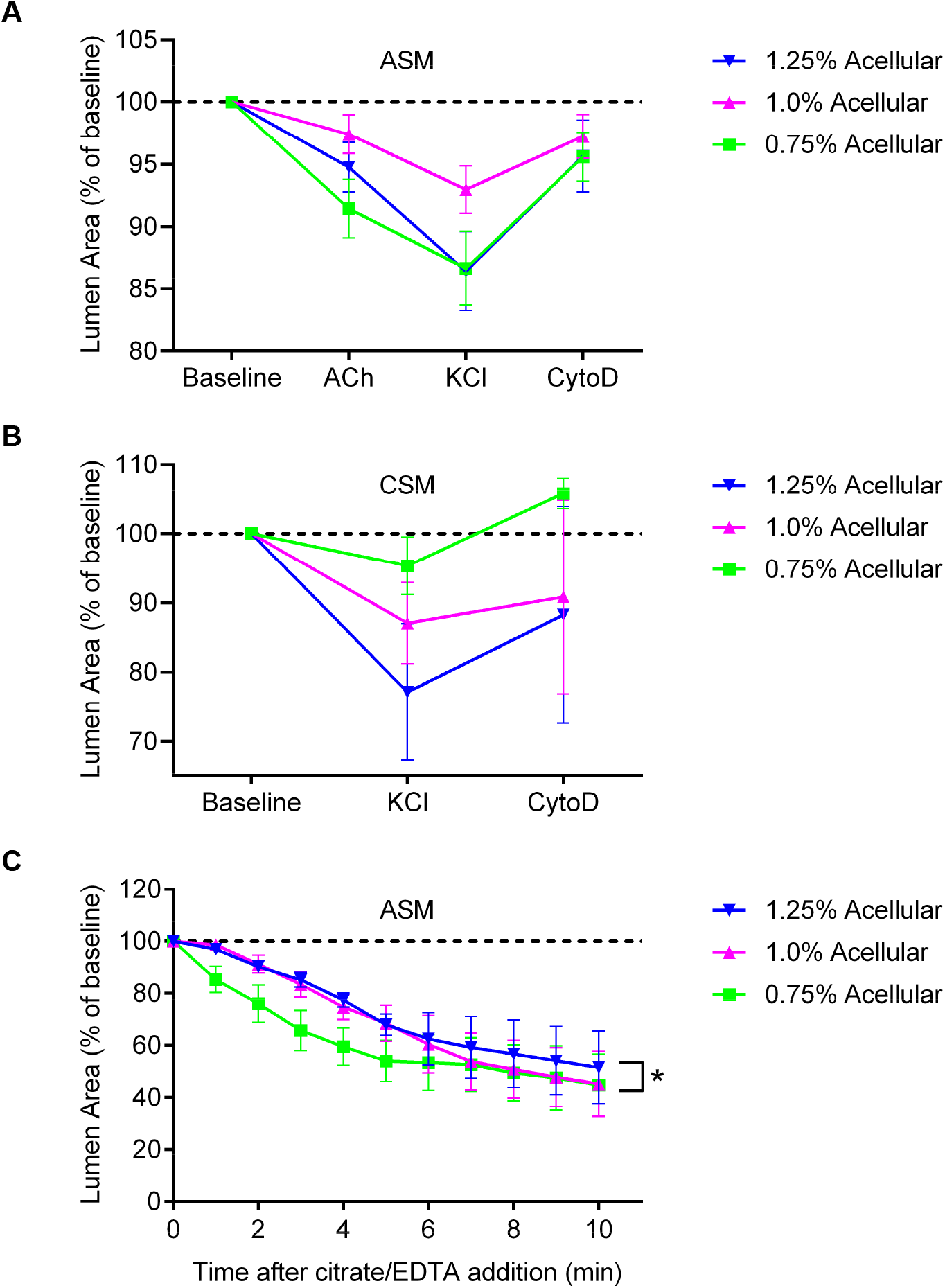
Contraction and relaxation of ASM and CSM sandwich designs. ASM sandwich constructs exhibit a drug- and stiffness-dependent contractile response (A; interaction p=0.5524, drug p<0.0001, stiffness p=0.0438, n=10), with mild contraction responses to acetylcholine, a stronger response to KCl, but only partial reversal of contraction by cytochalasin D. Surprisingly, the 1.0% acellular constructs exhibited significantly weaker response, despite being the median of the stiffness range. CSM also appeared to have similar contractile responses, but no significant differences were detectable due to high variation between samples (B; interaction p=0.8110, drug p=0.1788, stiffness p=0.2726, n≥4). Chelation-depolymerization of the alginate load in ASM sandwich constructs resulted in time- and stiffness-dependent reductions in lumen area (C; interaction p=0.9981, time p<0.0001, stiffness p<0.0074, n≥2), indicating that the muscle has substantial baseline tone, and that contraction can cause plastic deformation of the sandwich. * represents a significant stiffness effect.

The inability of cytochalasin D to completely reverse KCl contractions and expand lumen area above baseline levels was unexpected. This result presents two possibilities: 1) ASM and CSM in our model have little-to-no baseline contractile tone, or; 2) the sandwich can prevent ring collapse during tissue maturation but is insufficient to create an elastic load that can oppose strong or rapid contractions. To ascertain the relative degree of baseline tone versus strength of the load, ASM sandwich constructs were treated with a citrate/EDTA solution to chelate calcium and depolymerise the alginate, resulting in a sudden removal of the load opposing contraction. Figure 4C shows that load removal results in significant time (p<0.0001) and stiffness (p=0.0074) linked reductions in lumen area that far exceeds the magnitude of KCl contractions. These combined data indicate that the muscle exerts a significant degree of baseline contractile tone, and that incomplete recovery of lumen area with cytochalasin D is likely due to imperfect elasticity (i.e., plastic deformation) of the acellular structure.

### Spiderweb Development

To reduce plastic deformation of sandwich constructs during contraction, we further iterated on the tissue design to improve the ability of the acellular structure to behave like an elastic spring. This could not be feasibly achieved with a bioprinted acellular structure alone (data not shown). Thus, we designed a FDM 3D printed polycarbonate tissue holder to sit tightly within a ThinCert insert to create a rigid, non-compliant load against which the construct could develop tension. Six posts were precisely positioned in the holder, and hand-coded bioprinter toolpaths were created comprising circular ‘loops’ around the post positions. A constant direction of rotation, careful optimisation of loop length/width, and tight control of bioprinting speed and pressures were found to be necessary to minimise slack without closing the loops. Five layers of looping acellular structure was coded to encapsulate two layers of sandwich infill above and below the muscle bundle, which was reduced from 15 to 10 layers to balance muscle force relative to load strength and increase longevity in culture. Tissue holders were manually placed on top of the bioprinted structure immediately after printing, completing the spiderweb design and preparing the constructs for automated imaging (Fig 5A-B).

**Figure 5.**
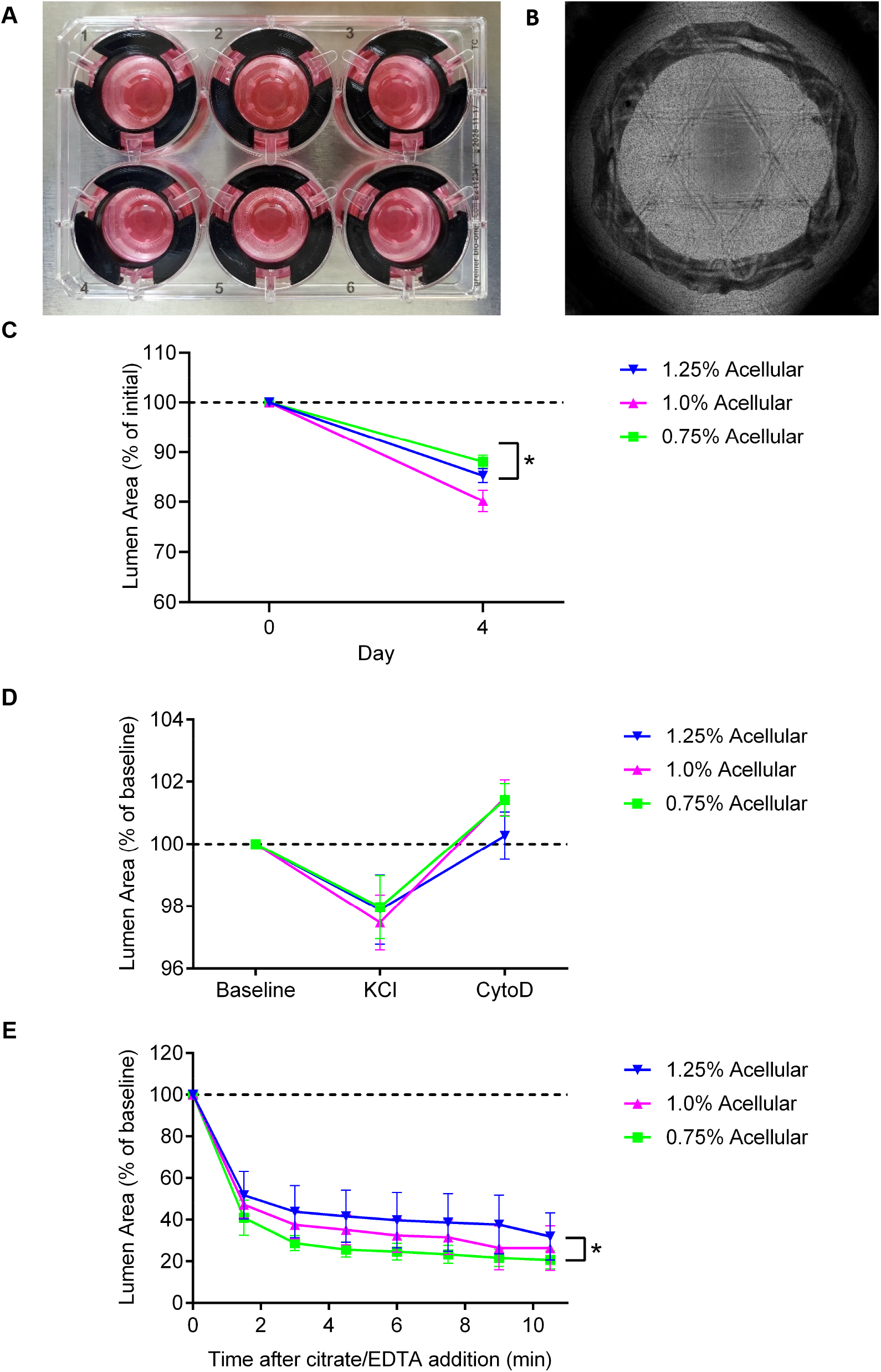
The spiderweb design presents a strong and elastic load opposing ASM contraction. The ASM ring within spiderweb designs is fixed in place by ThinCert centring rings, the tissue holder, and the taut acellular structure (A), enabling automated capture of high-resolution montage images for accurate lumen area assessment (B). Compaction of ASM spiderweb constructs followed a similar trajectory to ASM and CSM sandwich constructs, being modulated by both time and stiffness (C; interaction p=0.0036, day p<0.0001, stiffness p=0.0036, n≥16). The contractile response was drug-dependent (D; interaction p=0.7925, drug p<0.0001, stiffness p=0.7656, n≥5); modest reductions in lumen area were observed after KCl, while cytochalasin D completely reversed the contraction and enlarged lumen area above baseline levels. The chelation-depolymerization load removal resulted in a dramatic reduction in lumen area (E; interaction p>0.9999, time p<0.0001, stiffness p=0.0126, n≥3), indicating that the muscle in ASM spiderwebs exerts significant baseline tension, and that the spiderweb provides a strong elastic load that opposes the generated forces. * represents a significant stiffness effect.

Assessment of ASM spiderweb compaction at 4 days post printing (Fig 5C) revealed similar results to the ASM sandwich. There was significant reduction in lumen area over time (p<0.0001) and across the stiffness range (p=0.0036), although somewhat surprisingly the 1.0% acellular constructs showed modestly higher compaction.

### Spiderweb Contractile Function

Spiderweb constructs responded predictably to KCl and cytochalasin D with significant contraction and relaxation phases (Fig 5D; p<0.0001). However, in contrast to the sandwich design, spiderweb stiffness alone was unable to modulate contractile responses (p=0.7656), lumen area reductions to KCl were of a smaller magnitude, and lumen area increases to cytochalasin D rose above baseline levels. The small magnitude of spiderweb contractions relative to sandwich contractions could be attributed to reduced contractility and/or increased load. To clarify this question, alginate depolymerization experiments (Fig 5E) were performed revealing a strikingly rapid reduction in lumen area (p<0.0001) that was stiffness modulated (p=0.0126), with 0.75% acellular constructs showing the strongest response. Importantly, the contraction appeared to be considerably faster and more extensive than observed for sandwich constructs (Fig 4C). This suggests that the spiderweb muscle exerts a high degree of baseline contractile tone, and that the acellular load provides significant elasticity to oppose this force.

The dynamics of tension development in spiderweb designs was assessed by exposing a 0.75% acellular construct to KCl and cytochalasin D for 20 minutes each (Fig 6A). The contraction phase resulted in a steady reduction in lumen area that did not fully peak after 20 minutes. Cytochalasin D application resulted in a rapid and extensive reversal of contraction with lumen area ending up larger than baseline levels, indicating the spiderweb successfully generated a more effective spring than the sandwich.

**Figure 6.**
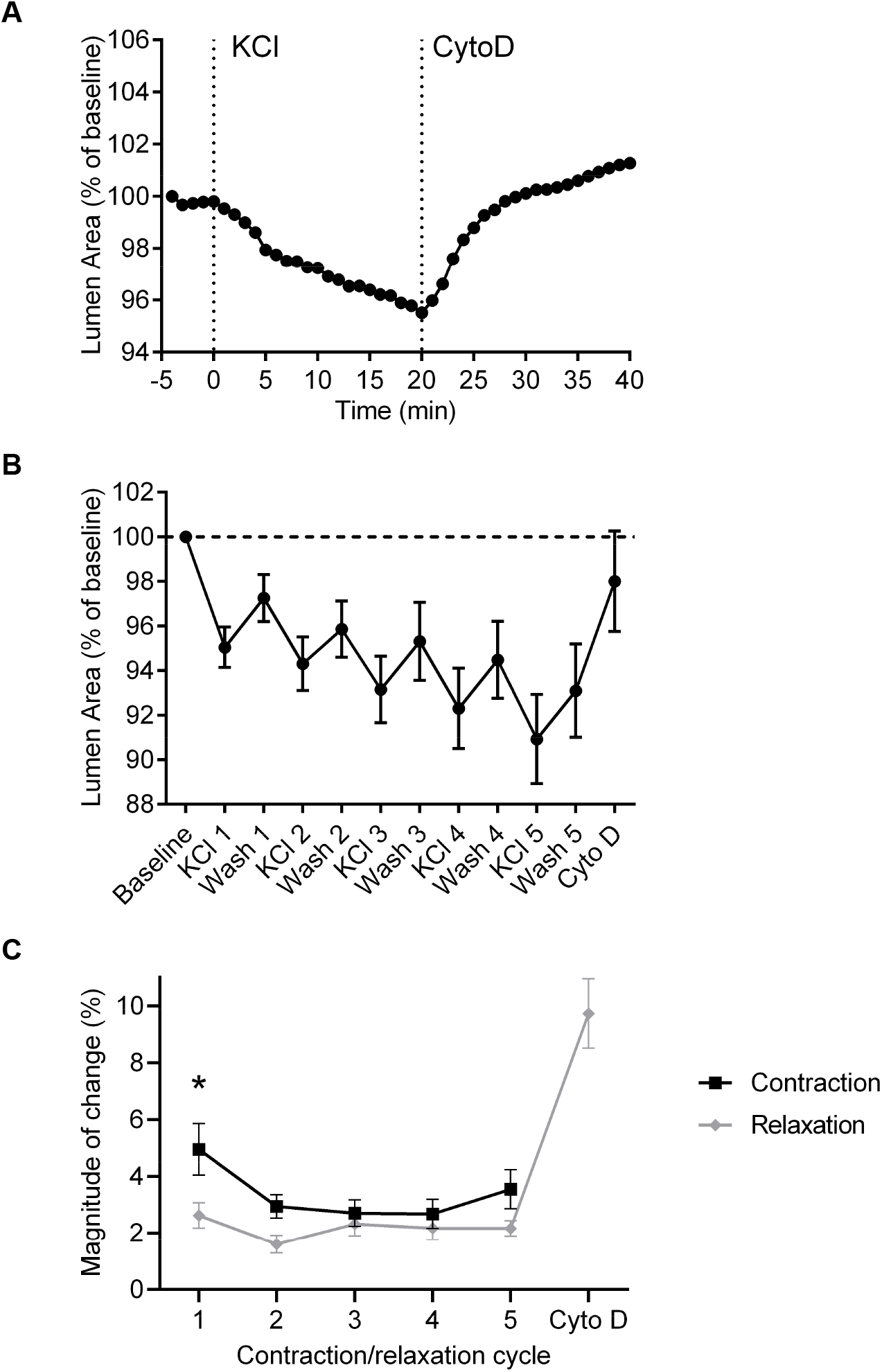
Dynamic properties of ASM spiderweb contractions. Tracking a single contraction (A) reveals that a maximal dose of KCl results in a steady reduction in lumen area that does not peak after 20 minutes. Cytochalasin D results in lumen areas larger than baseline, entirely reversing the contraction plus highlighting baseline tone. Repeated contraction/relaxation cycles cause a progressive reduction in lumen area (B), resulting from each contraction being larger than the subsequent relaxation (C; interaction p=0.3465, cycle # p=0.0314, contract vs relax p=0.0005, n=10). Cytochalasin D can partly reverse the cumulative reduction in lumen area, indicating contributions from increasing muscle tension and plastic deformation of the construct. The final cytochalasin D treatment is shown for reference but is excluded from the statistical analysis. * represents a significant difference between contraction and relaxation.

During repeat contraction/washout cycles, lumen areas of 0.75% acellular spiderweb constructs progressively decreased (Fig 6B). Analysis of each cycle (Fig 6C) revealed that the magnitude of contraction during the first cycle was significantly larger than subsequent cycles (p=0.0314), and that contraction was significantly larger than washout relaxation across the experimental time course (p=0.0005). A final treatment with cytochalasin D after the last cycle resulted in significant additional relaxation, indicating a progressive increase in baseline muscle tone. Since cytochalasin D could not fully return lumen area to baseline levels, this suggests that the spiderweb construct is still subject to plastic deformation when exposed to excessive contractile force.

## DISCUSSION

Creating physiologically relevant smooth muscle suitable for evaluating contraction remains a significant challenge for tissue engineering. This is particularly important within the context of the lung, where structure, function, and forces generated by ASM are inextricably linked. The recent description of a novel bioprinted experimental model of ASM (21) raises intriguing possibilities to create customised tissues that replicate different mechanical environments associated with development, health, and disease. In this study, we describe the development of innovative multi-material tissue designs that were enabled by the RX1 bioprinter’s ability to seamlessly switch between cellular and acellular bioinks within a single tissue engineered construct. Combined with a stiffness-tunable alginate-collagen-fibrinogen bioink, this allowed us to create widely different mechanical contexts for tissue engineered ASM. Importantly, we demonstrate that changes in the mechanical environment significantly modulate ASM contractile function.

### Structural Modulation of Tissue Engineered Muscle

Our study demonstrates the potent ability of tissue structure to drive a wide range of muscle function from the same cells and bioink components. Importantly, we highlight the pitfalls of not considering mechanical factors when engineering lung tissues that include a muscle component. Bare rings that represent muscle with an inappropriate mechanical load opposing contraction rapidly collapse and become non-contractile, even as cell viability and cellular morphology may remain qualitatively acceptable. Muscle formed in this manner may meet many metrics of experimental success but has no *in situ* physiological equivalent and would likely provide artifactual results for any experiment relating to contractile function.

The encapsulated sandwich and spiderweb designs represent a physiologically relevant mechanical load, and direct parallels can be drawn to the structure and function of different lung tissues. The mild mechanical load and resulting contractile dynamics offered by the sandwich design could be considered equivalent to a small non-cartilaginous airway. In this context, the mild load allows a relatively high degree of muscle shortening during contraction, and contraction/relaxation plastic deformation of the structure may simulate airway closure. Further, the relatively slow rate of shortening observed when the mechanical load is removed by depolymerising alginate with citrate/EDTA is comparable to the relatively low baseline tone seen in small airway smooth muscle. In contrast, the more robust load provided by the spiderweb design appears more akin to large cartilaginous airways. The magnitude of muscle shortening during KCl contractions is restricted by the load, and the rapid shortening upon load removal is strikingly similar to the shortening that occurs during dissection of highly tonic tracheal smooth muscle.

Within each tissue design, stiffness modulation of the structural component produced somewhat inconsistent results. The predicted weaker loads produced by the 0.75% acellular bioink allow greater sandwich compaction during ASM tissue formation, yet spiderweb compaction was significantly greater in 1.0% acellular ASM constructs, and no statistically significant difference was observed in CSM. Load removal experiments exhibited more consistency; in both sandwich and spiderweb tissues, 0.75% acellular constructs exhibited faster and more extensive shortening during load removal, as expected from the reduced quantity/density of alginate that needs to depolymerise. Contractile responses were also diverse in the different tissues and structures trialled; ASM sandwiches had the smallest contraction in 1.0% acellular tissues, CSM sandwiches trended towards larger contractions with increasing stiffness, and ASM spiderwebs contracted similarly irrespective of acellular stiffness. A key confounding factor that possibly contributed to inconsistencies in contractile function is that our results were normalised to a baseline lumen area recorded immediately prior to contraction experiments. This is directly equivalent to standard protocols that set baseline length (28) or optimal passive force (29) prior to *ex vivo* smooth muscle strip experiments. However, adopting this approach in our model masks the effect of varying levels of muscle maturation and tissue compaction preceding evaluation of contractile responses, which may represent different degrees of tissue remodelling versus baseline tone development. Moving forward, it will be essential to image every construct immediately after printing and before contraction experiments to account for this variable.

It is also possible that the effects of differing acellular load stiffness are below the sensitivity of instruments used to capture contractile responses. Lumen area changes in spiderwebs are notably small and may be subject to noise despite our rigorous imaging and measurement protocols. We expect that clearer patterns would emerge with a wider range of acellular stiffnesses within each tissue design. In this study we used acellular SLG-20 alginate concentrations from 0.75% to 1.25%, representing the lowest and highest bioink viscosities printable with high fidelity and near-perfect reliability. Based on printing pressures (Table 1) this range covers an approximate doubling of bioink viscosity and resultant fibre stiffness. However, to better highlight stiffness modulation of contraction, an expanded concentration range of 0.5 to 1.5% SLG-20 is entirely feasible (albeit with a lower success rate of producing high fidelity constructs) and would be expected to produce more than a three-fold difference in fibre stiffness between the softest and stiffest constructs. Finally, this foundational design study primarily considers muscle function in terms of physical shortening, which is a relatively narrow view of the many potential factors modulated by tissue mechanics. Future studies will examine other experimental outcomes with a wider dynamic range of measurements including contractile vs synthetic phenotype, calcium handling, proliferation, and ECM remodelling.

### Contrasts to Other Tissue Engineering Approaches

The bare ring, sandwich, and spiderweb designs developed in this study offer striking similarities and disparities to comparable hydrogel models of tissue engineered ASM. Notably, a foundational 3D bioprinting paper by Dickman *et al* used an alginate-collagen bioink (AGC-10) to create primary human ASM muscle rings that compacted without collapsing [see Ref (21) Fig 2]. Our inability to replicate similar data may be related to distinct properties of AGC-10, or attributable to use of ASM cells with significantly different mechanical and contractile properties. Indeed, discussions with the authors revealed that muscle rings made with some ASM cell samples did collapse, and these tissues were excluded from their experiments. Wide variation in primary ASM cell samples is normal and expected, but such an approach may bias cell selection towards low-contractile cells incapable of generating high amounts of force. This highlights a key benefit of our designs; the acellular layers generate a significant load opposing cellular tension, allowing both weakly and highly contractile cells to be used in identical conditions without fear of collapse.

The ASM microtissue model (18) provides an excellent point of comparison since the studies used the same ASM cell line and key contractile experiments were performed with similar protocols. Indeed, several physiological aspects of the sandwich and spiderweb constructs were remarkably similar to microtissues, including actin morphology, the relative magnitude of ACh and KCl contractions, and time courses for maximal contraction and relaxation. Yet, a marked point of difference between the microtissue model and this study lies in the dynamics of multiple contraction and washout relaxation cycles in spiderweb constructs [see Ref (18) Fig 4E]. In both experimental models the first contraction was greater than subsequent contractions, but in spiderweb designs each contraction was of greater magnitude than the subsequent washout relaxation, resulting in a progressive decrease in lumen area with each cycle. The incomplete relaxation after the final cytochalasin D treatment indicates that this phenomenon derives in-part from plastic deformation of the spiderweb throughout the experiment. However, the large magnitude of cytochalasin D relaxation suggests that the washout step alone is incapable of fully relaxing the muscle, and that active actinomyosin crossbridge cycling and/or passive cytoskeletal tension accumulates with successive contractions.

The reason for this contrasting relaxation response is unclear, although key technical differences between the models may contribute. Spiderweb muscle bundles are significantly larger than microtissues with a greater diffusion barrier that potentially increases washout times. However, single-contraction time courses were highly comparable between the models indicating this is not a major factor. The inclusion of supporting fibroblasts in ASM microtissues is noteworthy, although it seems doubtful that fibroblasts would modulate muscle relaxation in this manner. More likely, limitations on microtissue lifespan mean they were cultured for shorter time periods than spiderwebs (typically 3 days versus 5 days) and had less exposure to low serum media (2 hours versus ≥24 hours), each of which may significantly modulate muscle relaxation. Finally, the silicone cantilevers of the microtissue model behave like perfect springs, with virtually no hysteresis nor plastic deformation, which may actively pull on the tissues to facilitate lengthening during relaxation versus the less elastic spiderweb design. Examining other features relating to relaxation, including the use of alternative relaxant agents (β_2_ agonists, adenylate cyclase activators, rho kinase inhibitors), tracking cytoplasmic calcium, and characterising cytoskeletal morphology will be essential to develop a complete understanding of this differential relaxation behaviour.

In terms of practicalities, our bioprinted designs offer significant benefits over microtissues and other models based on deformable silicone moulds. All aspects of the sandwich and spiderweb designs are far easier and less resource intensive to produce than microtissue and other silicone substrates, tissue mechanical properties can be altered ‘on the fly’ without changing silicone manufacturing processes, and the entire bioprinting process requires only off-the-shelf equipment and reagents. The bioprinting process can be completed by a moderately skilled operator and does not require laborious and highly skilled preparation of substrates including degassing of wells, centrifugation, or substrate ‘scraping’ steps. Importantly, bioprinting may be more suitable to obtain reliable results from rare or expensive cell samples. While large quantities of cells are required for each bioprinting run (≥5×10^6^ cells), virtually all of the cells can be incorporated into constructs. Only 1×10^6^ cells are required for manufacturing microtissues, but it is estimated that less than 5×10^4^ cells end up within constructs due to manufacturing losses associated with centrifugation and scraping. Finally, bioprinted tissues last significantly longer in culture than microtissues and automated measurement protocols are easier to establish, making them suitable for a much wider range of experiments.

### Other Limitations and Opportunities for Deconstructing the Lung

The narrow range of alginate types and concentrations that generated functional smooth muscle tissue was surprising. We had initially expected to create a bioink based on RGD-VLVG where muscle bundle stiffness could be widely modulated by addition of stiffer alginates. However, alginate content needed to be kept relatively low, and inclusion of higher molecular weight alginates – even at concentrations as a low as 0.1% – completely inhibited tissue formation. This may be consistent with previous studies that show high native alginate concentrations impact cell function due to poor diffusivity and porosity (22, 23), and that alginate chain length specifically regulates cell spreading (25). In this context, a bioink comprised of ultra low molecular weight alginate generated by alginate lyase treatment (24) may provide greater scope for modulating the stiffness of the muscle bundle, since alginate concentration could likely be increased without inhibiting cell spreading. This approach was not pursued in this study; the ability of the sandwich and spiderweb designs to completely uncouple the cellular and acellular components provided significant freedom for construct optimisation. Nevertheless, the inability to use alginate to stiffen the muscle bundle may reduce the capacity to simulate some aspects of fibrosis, and alternative approaches such as matrix crosslinking (30, 31), inclusion of fibroblasts (18), or treatment with mediators of fibrosis (21) may be more fruitful.

In contrast to the rigid limits on alginate composition, the wide range of ECM protein content achievable within the bioinks was impressive. Inclusion of small amounts of collagen I significantly improved structural integrity immediately after printing, likely due to collagen thermal gelling contributing to layer adhesion. Collagen I also improved tissue formation versus RGD-alginate alone, consistent with its well-known role for improving biocompatibility in cell culture and tissue engineering applications. Fibrinogen inclusion (with post-print conversion to fibrin) was not inherently required to create contractile ASM, but was beneficial for improving long-term structural integrity of constructs. This is expected from the excellent mechanical properties of fibrin/collagen co-gels (32), the resistance of fibrin to degradation by proteases normally secreted by smooth muscle (33–35), and the absence of plasminogen in our culture system. Intriguingly, the upper limit of collagen and fibrinogen concentrations was restricted only by practicalities of stock solution concentrations and the quantity that could be physically mixed into the bioink while still maintaining osmolarity and pH. This raises the possibility that collagen and fibrinogen could be, in-part or completely, substituted by alternate ECM proteins including decellularized ECM (36, 37) to test the effects of different ECM milieu on muscle function. Usage of dECM may be particularly beneficial, as the inclusion of dECM in bioprinted intestinal smooth muscle constructs was shown to improve contractile function (21). Moreover, the nature of the sandwich and spiderweb designs mean that these ECM substitutions can be made freely without concern for interfering with muscle loading, creating a powerful tool to study how ECM derangement interacts with tissue mechanics.

### Summary

We have developed a physiologically relevant bioprinted model of ASM that enables extensive manipulation of tissue mechanics. The combination of a stiffness-tunable bioink, partnered with a microfluidic-based multi-material bioprinter than enables the novel sandwich and spiderweb designs, allows us to interrogate the effect of lung structure on ASM contraction more readily than any other tissue engineered model. Future studies examining a broader range of stiffness environments, alternative ECM components, and simulated fibrosis by including fibroblasts or through ECM crosslinking, will provide valuable insights into lung development, health, and disease.

## Supporting information

STL and FDM Print Files

## SUPPLEMENTARY FILES

Supplementary files are available as a private link for review at https://figshare.com/s/a36437e31e92fbb4fb7e

The linked zip file contains:

Bare Ring.STL (STL file for 10 mm bare rings)

Sandwich.STL (STL file for sandwich design with 13 mm frame and 10 mm muscle ring)

Tissue Holder Thincert v4 22.8.STL (STL file for spiderweb tissue holder)

Tissue Holder Retraction Count Buster.STL (STL file for sacrificial filament usage)

UM PC for Tissue Holder.CURAPROFILE (printing profile for spiderweb tissue holder)

UMS3_Tissue Holder Thincert v4 22.8 (Cura 4 Project file for tissue holder for Ultimaker S3)

Thincert Centring Holder v3.STL (STL file for Thincert centring ring)

BASF Ultrafuse PET for Centring Holder.CURAPROFILE (printing profile for Thincert Centring Ring)

UMS3_Thincert Centring Holder v3 (Cura 4 Project file for Thincert centring ring)

## ACKNOWLEDGEMENTS

The authors would like to thank Samuel Wadsworth, Erin Bedford and Simon Beyer from Aspect Biosystems for helpful discussions and technical support relating to bioink design strategies, 3D design, and hand-coding bioprinter toolpaths. We thank Dr Andrew Halayko (Department of Physiology & Pathophysiology, University of Manitoba) for supplying the D12 ASM cell line.

## AUTHOR CONTRIBUTIONS

Jeffery Osagie edited the manuscript, designed experiments, and performed data/statistical analysis, cell culture and contraction assays

Sanjana Syeda edited the manuscript, designed experiments, and performed data/statistical analysis, cell culture and contraction assays

Emily Turner-Brannen edited the manuscript, analysed data, and performed bioprinting, cell culture and contraction assays

Michelle Guimond performed 3D FDM design

Lumiere Parrenas tested 3D FDM designs, and performed cell culture and contraction assays

Ahsen Haroon tested 3D FDM designs, and performed cell culture and contraction assays

Philip Imasuen tested 3D FDM designs, and performed cell culture and contraction assays

Adrian R West conceived the study, wrote the manuscript, designed/supervised all aspects of experimentation, and performed data/statistical analysis, 3D design, FDM 3D printing and bioprinting

